# A configural model of expert judgement as a preliminary epidemiological study of injury problems: An application to drowning

**DOI:** 10.1101/517300

**Authors:** Damian Morgan, Joan Ozanne-Smith

## Abstract

Robust epidemiological studies identifying determinants of negative health outcomes require significant research effort. Expert judgement is proposed as an efficient alternative or preliminary research design for risk factor identification associated with unintentional injury. This proposition was tested in a multi-factorial balanced experimental design using specialist judges (N=18), lifeguards and surfers, to assess the risk contribution to drowning for swimming ability, surf bathing experience, and wave height. All factors provided unique contributions to drowning risk (*p*<.001). An interaction (*p*=.02) indicated that occasional surf bathers face a proportionally increased risk of drowning at increased wave heights relative to experienced surf bathers. Although findings were limited by strict criteria, and no gold standard comparison data were available, the study provides new evidence on causal risk factors for a drowning scenario. Countermeasures based on these factors are proposed. Further application of the method may assist in developing new interventions to reduce unintentional injury.

## Introduction

Analytic epidemiological studies test for the association of determinants with a negative health outcome to support a theory of causality. Identified causal risk factors may then be modified to improve health outcomes. Epidemiological research designs provide robust evidence through observing candidate risk factors in the natural course of events. Means of potential risk factor identification and specification include anecdotal evidence, case reports or cross-sectional surveys.

A significant challenge for observational epidemiological studies concerns the control of confounding factors [1]. Elimination of competing explanations for study findings often requires substantial study sizes. Accurate measurement of exposure to risk factors is complex as is distinguishing factor causation from statistical association. Without attention to these details, epidemiologic studies provide little substantial knowledge gain.

Uncertainly about the derived health benefits leads to difficulty in justifying study costs and may partially explain the lack of epidemiological studies for many significant injury problems. To provide evidence on potential health gains, cost-effective preliminary studies with acceptable internal validity and generalisability may guide later epidemiological research. Such studies should replicate more rigorous epidemiological designs with respect to risk factor identification and assessment. The study reported here tests and evaluates a proposed method based on expert opinion for potential causal injury risk factor identification and risk quantification as applied to unintentional drowning in an Australian surf bather population, where surf bather drowning is a relative rare event relative to bather numbers [2].

## Drowning as a global health problem

Globally, drowning accounts for more than 370,000 deaths each year [3]. The causes of drowning are complex and risk factors vary by geographic location, physical environmental features, activity, water entry mechanism, weather and water conditions, supervision, and personal characteristics. Therefore, studies of causal risk factors should be restricted to clearly defined circumstances.

In most drowning scenarios, including surf bather drowning, scant evidence exists on causal risk factors. Surf bathing at wave-dominated beaches attracts local residents and is commonly depicted in tourist brochures luring visitors to warm and exotic beach locations [4]. Despite dedicated beach patrols and lifeguards supervising bathers, Australia’s annual coastal drowning rate remains 0.14 swimmers and waders per 100,000 resident population [5]. Several causal drowning risk factors have been proposed for surf bathers, including rip currents, alcohol, tourists, and onset of medical conditions [6]. Yet no rigorous research studies confirm that these or other candidate causal risk factors place bathers at relatively higher drowning risk.

This study aimed to test the capacity of experts to identify and assess the roles played by putative causal risk factors in surf bather drowning.

## Judgement and risk assessment

Expert judgement provides a recognised method for gathering evidence where traditional scientific methods are impractical [7-8], such as assigning risk probabilities to assess certain environmental hazards [9-10]. Such risk assessments may be biased by judge overconfidence, inaccuracy, or insufficient or irrelevant judge expertise [11-12]. Given these and other potential limitations, experiments based on subjective judgements require control of recognised potential bias and careful selection of judges, thus limiting generalisability of the findings.

Early judgement studies sought to model the decision-making process and assess the applicability of outcomes to the *true state* [13-15]. Linear modelling has matched the process used by judges where variables are assigned weights, with the sum used to determine the outcome likelihood. In reality, judges may follow a configural rather than a linear process by assigning values to predictor variables based on weights of other predictor variables [14]. Related statistical techniques such as analysis-of-variance (ANOVA) can account for configurality within judgements (predictor variable interactions).

## Method

Fig 1 presents the study design overview. Based on a repeated-measures multi-factorial experimental design, two separate groups of surf bathing *specialists*—lifeguards and surfers—were recruited to judge the contribution of putative causal factors to surf bather drowning risk.

**Fig 1.**
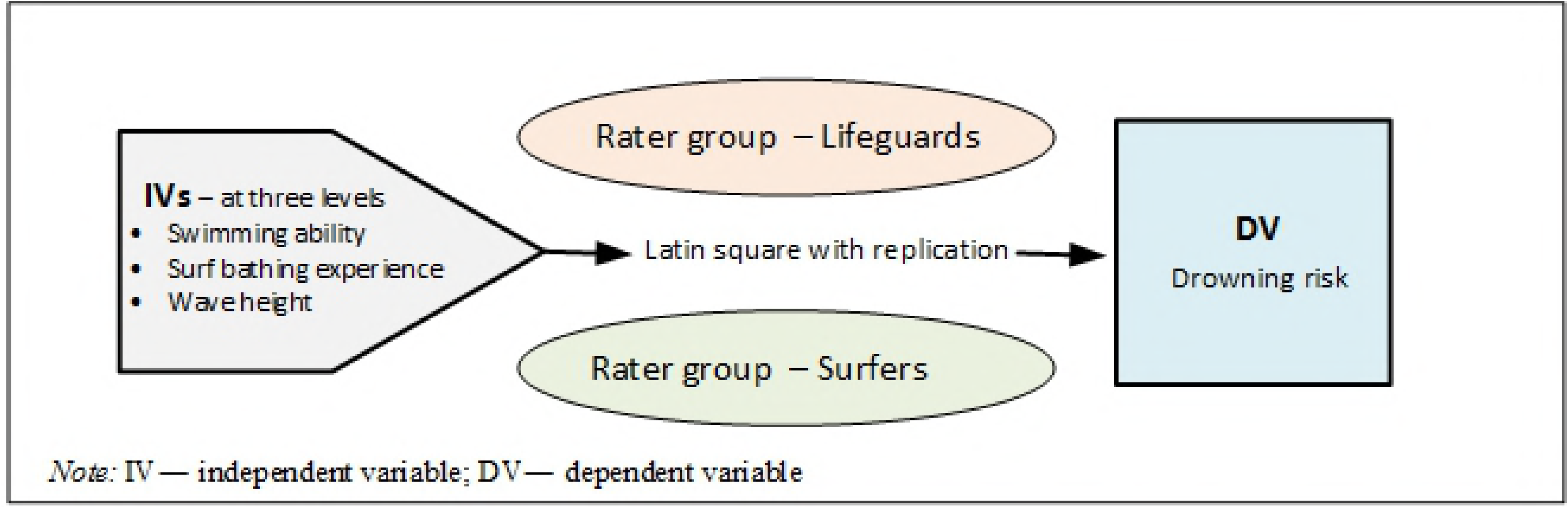
Study design overview.

### Independent variables and factor levels

A preliminary study eliciting water safety expert knowledge, using a nominal group technique, reported swimming ability in surf conditions, awareness of surf hazards, and prevailing surf conditions as the top ranked factors affecting the probability of surf bather drowning [16]. From this, three independent (predictor) variables (IVs) were specified as putative drowning risk factors: *swimming ability, surf bathing experience*, and *wave height*. IVs were set at three fixed levels representing ordinal scales, restricting inferences to the specified IV levels [17]. Factor levels were distinguished by a mix of qualitative descriptions and quantitative measures (S1 Table). Levels were ordered from (presumed) lowest to highest drowning risk contribution, assuming other factors are absent (e.g., alcohol), remain equal or constant (e.g., tide level or health status). For each factor, the median risk level was anchored at averages found for Australian beaches or surf bather populations [18-19].

### Dependent (criterion) variable

A scale measurement was required for surf bather drowning risk. Piloting revealed judge preference for the term ‘getting into difficulty’ as a proxy scale measure of drowning risk. This scale appraised the likelihood of bathers reaching their limit to cope with surf conditions based on their swimming ability and surf bathing experience and logically, this situation is a precursor to drowning. This scale was used as the proxy drowning risk measure for the study.

The dependent (criterion) variable (DV) used an 11-point scale to record the perceived chance of getting into difficultly while bathing—0% to 100% [20]. Descriptive terms below the scale qualified associated percentage ranges for getting into difficulty: No; low; moderate; high chance; and certain (S1 Table).

### Hypotheses

Three hypotheses (Hs) were specified:

H1: *The IVs*—*swimming ability, surf bathing experience, and wave height*—*will each produce an effect on the DV*—*chance of getting into difficulty in the water*.

Each of the three scale items was expected to be associated with varying levels of surf bather drowning risk in a systematic *risk* order providing the rational for H2.

H2: *The order of levels within each IV is associated systematically with surf bather drowning risk*.

Null-hypotheses tested for differences between IVs levels and expected direction.

Intuitively, and consistent with a configural judgement approach, the drowning risk for levels of one IV would be expected to be influenced by the other two IV levels. Therefore, H3 anticipated that an effect produced on the DV by one level of an IV is dependent on levels of other IVs [21].

H3: *Three first order interactions and one second order interaction among the IVs will produce an effect on the DV*.

H3 generated four null-hypotheses. The direction of interaction effects between IV levels were investigated *a priori* but not as specified hypotheses.

### Instrument and design

Ethical approval was granted by the Monash University Standing Committee on Ethics in Research Involving Humans. The experiment was administered using a self-completed questionnaire. Personal data comprised surf bathing experience and currency, surf-activity proficiency, lifesaver/lifeguard and rescue experience and demographic details. To reduce potential influence of other possible risk factors on judgements, an instruction page outlined the general scenario for drowning risk exposure including bathing at the outer wave breaking zone (S2 Table). Following this, 27 vignettes, on separate pages, provided a combination of IV levels and the DV drowning risk scale. Two sets of questionnaires were produced; P1 for time-period 1 and P2 for time-period 2. Respondents were instructed not to refer back to their previous ratings when rating new scenarios.

### Vignettes and ordering procedure

The three IVs at three levels resulted in 27 unique combinations (cells) for rating the DV. Each cell was presented as a three paragraph vignette personalised with a gendered name (S2 Table).

The order of IVs can affect the DV score due to participant practice, fatigue or becoming *wise* to the experiment [21]. To counterbalance carry-over (order) effects, a Latin square type arrangement was used to distribute carry-over effects systematically across cells [17, 22]. It was anticipated that statistical analysis would account for carry-over effects within error terms. The sequence of IVs and subject gender within each vignette was also systematically varied.

A repeat square for P2 provided supplemental data to assess the reliability of responses between P1 and P2. Effectively, the use of the repeat square provided two cell ratings per judge under different order conditions [23].

### Population and sampling procedure

Expertise encompasses skills and knowledge and experts may be identified for specific areas from characteristics including capabilities, achievements, qualifications, peer recognition, specialisation or years of performance [24-25]. Two populations were considered expert in surf bathing activities, professional lifeguards and proficient surfers, allowing comparison of judgements across specialist populations. Through bather supervision, lifeguards have direct experience of bathers getting into difficulty and hold recognised proficiency in surf bathing activities. Surfer expertise encompasses the necessary skills and experience to negotiate typical and atypical surf conditions.

### Sample size and statistical power

Stevens [26] provides required sample sizes for single group repeated-measures ANOVA. To obtain 80 percent power, assuming an average correlation of DV measures being 0.5, three treatments (IVs), alpha level of 0.05, and large effect size, 8-14 repeated-measures are required. The target sample size was nine lifeguard and nine surfer judges. Statistical power may be increased through pooling of results where no statistical differences are found between the specialist groups or time periods.

### Selection and study participation

Participants were selected using a convenience sampling procedure following a snowball-like process. Some participants were known to each other. All questionnaires were completed in DM’s presence. Following instruction, 18 participants completed the first set of 27 vignettes (each in a unique order) followed by a 30 minute break. Demographic information and the second set (repeat square) were then completed.

## Data analysis

Introduced bias was firstly assessed for vignette gender, time-period effect, and specialist type. Following this, factorial repeated-measures three-way ANOVA established simple main effects and error terms for the IVs (H1). Statistical differences for IV levels and direction were then assessed (H2). Planned polynomial contrasts between IVs specified interactions (H3).

The ANOVA results follow the order of hypotheses. Where an interaction between IVs was found, simple pairwise comparisons of means were examined *post-hoc* using the middle IV level (median risk level) as the reference group [27].

Preliminary data analyses revealed a wording error for Vignette 7—one IV factor level being incorrect. This resulted in estimated V7 DV scores for each judge being made by interpolation [28]. This took into account mean score patterns for corresponding vignette levels and applied differences to individual V7 scores [29]. The procedure maintained existing order effects and parallel risk assessments particular to each judge. Specifically, the score for V7 (with same procedure followed for the repeat square) was calculated as equal to: original score (V7) less mean difference between V9 and V8 less mean difference between V17 and V16. This resulted in 3 of 18 judges’ scores for P1 being negative (−1.2, −0.2, −0.2) and 1 judge for P2 (−0.5). These four negative scores were converted to zero. The face validity for the derived V7 mean score was confirmed by comparison with the *closest* cell levels. Remaining reporting treats the derived estimates for V7 as the true scores.

Data were entered on the spreadsheet and analysed using statistical software [30] with alpha level *p*<0.05. The DV results were entered as scores (0-10) corresponding to percentage indicated. Normality of distributions was assessed for each vignette visually and by reference to significance tests for skewness and kurtosis z-scores (*p*<0.05). Due to small sample sizes and potential non-normal distributions, non-parametric tests (exact significance) were used for preliminary subgroup comparisons.

Differences between specialist groups on demographic and beach behaviour were determined by Mann-Whitney [*U*] tests or chi-square [χ ^2^]) with corresponding effect size calculated manually for significant results. To test for vignette name gender effects, P1 and P2 data were combined for each of the 27 vignettes (each being rated 36 times, twice by a single judge) with differences assessed by Mann-Whitney (*U*) tests.

Reliability of judges’ scores over P1 and P2 was assessed by the Wilcoxon signed-rank test. Here, significant results at *p<*0.05 would be expected as the cumulative (family-wise) Type 1 error rate across 27 tests increased the likelihood of false positives. Bonferroni correction set a significant alpha level at *p<*0.002 [31]. DV ratings for each vignette were grouped for P1 and P2 to test for differences between specialist groups using Mann-Whitney (*U*) tests.

Following preliminary assessment, a repeated-measures three-way ANOVA was run on SPSS using the general linear program. The assumption of sphericity was assessed by Mauchly’s test. Greenhouse-Geisser estimate, correcting for degrees of freedom, was used where the assumption of sphericity was not met. *A priori* polynomial contrasts were specified for each factor to test for presumed IV factor order (in linear or quadratic form) with results reported where significant. Tests of differences between estimated marginal means for IV factor levels (i.e., the unweighted mean that controls for potential confounding from other IVs), three first-order interactions, and one second-order interaction, applied the Bonferroni correction. Partial eta squared (partial *η*^2^), which explains the proportion of variation unique to a variable not explained by other variables, was used to estimate effect sizes [27] with 95% CI calculated from SPSS syntax files from Smithson cited in [21].

Eta squared (*η*^2^) was calculated manually. This measure can be interpreted in a similar way to *R*^*2*^, being an additive portion of the total variance in the DV explained by the IVs and interactions, provided the design is balanced by an equal group size for each cell [21-22]. These tests for effect size were based on the sample results without correction for population estimates [27]. Figures were prepared manually using the Excel program [32].

## Results

### Specialist profiles

The specialist groups (lifeguards and surfers) had similar demographics and beach experience confirmed by non-significant differences on statistical tests (not reported). Surfers had higher frequency of beach visits in the previous 12 months (*U*=18.5, *p=*0.05). All judges had extensive experience in surf bathing (10 to 30 years). Most had high participation rates in the last 12 months and experience in 3 m waves. All judges except one rated themselves as proficient or expert in surf bathing.

All lifeguards held surf-related and first aid qualifications and had completed rescues, six having performed cardiopulmonary resuscitation. Almost half the surfer specialists held swimming-related qualifications, first aid certification or experience in performing rescues. A statistically significant difference was found for the average number of rescues performed by surfers and lifeguards (respective means 2.3 and 270.6, *U*<0.01, *p*<0.01, *r*= −1.2).

### Vignette gender

The DV mean rating (chance of getting into difficulty in the water) was higher for female vignette subjects compared to males for 17 vignette scenarios (63%); lower for 10 (37%). Mann-Whitney (*U*) tests were not significant for any vignette gender differences (*p*<0.05) so this variable was not treated as a factor in further analysis.

### Reliability of vignette DV ratings between P1 and P2

Vignette DV mean ratings between P1 and P2 (N=36) were 13 (48%) higher cell means for P1, 12 (44%) higher cell means for P2, and 2 (7%) identical means. As Wilcoxon sign-rank test identified no significant differences (*p*<0.002) P1 and P2 ratings were considered statistically to be from the same populations, providing justification for pooling cell ratings.

### Comparison of specialist groups DV ratings

Table 1 presents judges’ mean vignette ratings for P1, P2 and overall. Although vignette ratings varied (mean 4.0-6.9), when grouped the pattern for surfers and lifeguards were similar. Overall mean differences between specialist group ratings were not significant for P1, P2, or combined periods (P1: *U*=35.0, *p*=0.65, *r*= −0.11. P2: *U*=27.0, *p*=0.25, *r*= −0.28, combined: *U*=30.5, *p*=0.40, *r*= −0.21).

**Table 1.**
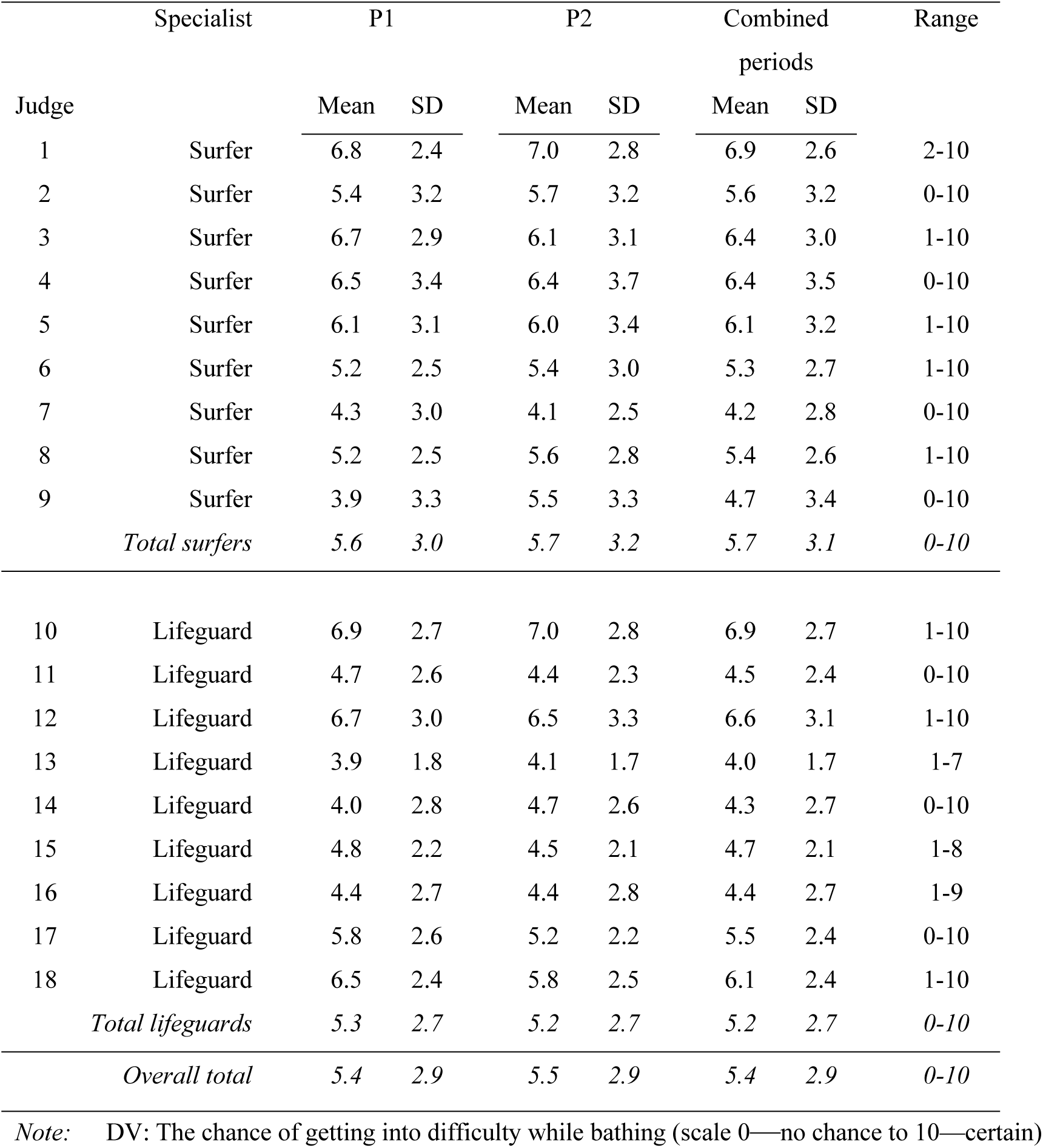
Specialist judges mean ratings of the DV for 27 vignette scenarios.

| Judge | Specialist | P1 |  | P2 |  | Combined periods |  | Range |
| --- | --- | --- | --- | --- | --- | --- | --- | --- |
|  |  | Mean | SD | Mean | SD | Mean | SD |  |
| 1 | Surfer | 6.8 | 2.4 | 7.0 | 2.8 | 6.9 | 2.6 | 2-10 |
| 2 | Surfer | 5.4 | 3.2 | 5.7 | 3.2 | 5.6 | 3.2 | 0-10 |
| 3 | Surfer | 6.7 | 2.9 | 6.1 | 3.1 | 6.4 | 3.0 | 1-10 |
| 4 | Surfer | 6.5 | 3.4 | 6.4 | 3.7 | 6.4 | 3.5 | 0-10 |
| 5 | Surfer | 6.1 | 3.1 | 6.0 | 3.4 | 6.1 | 3.2 | 1-10 |
| 6 | Surfer | 5.2 | 2.5 | 5.4 | 3.0 | 5.3 | 2.7 | 1-10 |
| 7 | Surfer | 4.3 | 3.0 | 4.1 | 2.5 | 4.2 | 2.8 | 0-10 |
| 8 | Surfer | 5.2 | 2.5 | 5.6 | 2.8 | 5.4 | 2.6 | 1-10 |
| 9 | Surfer | 3.9 | 3.3 | 5.5 | 3.3 | 4.7 | 3.4 | 0-10 |
|  | *Total surfers* | *5.6* | *3.0* | *5.7* | *3.2* | *5.7* | *3.1* | *0-10* |
| 10 | Lifeguard | 6.9 | 2.7 | 7.0 | 2.8 | 6.9 | 2.7 | 1-10 |
| 11 | Lifeguard | 4.7 | 2.6 | 4.4 | 2.3 | 4.5 | 2.4 | 0-10 |
| 12 | Lifeguard | 6.7 | 3.0 | 6.5 | 3.3 | 6.6 | 3.1 | 1-10 |
| 13 | Lifeguard | 3.9 | 1.8 | 4.1 | 1.7 | 4.0 | 1.7 | 1-7 |
| 14 | Lifeguard | 4.0 | 2.8 | 4.7 | 2.6 | 4.3 | 2.7 | 0-10 |
| 15 | Lifeguard | 4.8 | 2.2 | 4.5 | 2.1 | 4.7 | 2.1 | 1-8 |
| 16 | Lifeguard | 4.4 | 2.7 | 4.4 | 2.8 | 4.4 | 2.7 | 1-9 |
| 17 | Lifeguard | 5.8 | 2.6 | 5.2 | 2.2 | 5.5 | 2.4 | 0-10 |
| 18 | Lifeguard | 6.5 | 2.4 | 5.8 | 2.5 | 6.1 | 2.4 | 1-10 |
|  | *Total lifeguards* | *5.3* | *2.7* | *5.2* | *2.7* | *5.2* | *2.7* | *0-10* |
|  | *Overall total* | *5.4* | *2.9* | *5.5* | *2.9* | *5.4* | *2.9* | *0-10* |

*Note:* DV: The chance of getting into difficulty while bathing (scale 0—no chance to 10—certain)

DV rating patterns for individual vignette cells by specialist group provide further insight (Table 2). Table 2 shows broad similarity between surfer and lifeguard DV patterns across vignette cells. Upper and lower DV ratings for individual cells ranged substantially, partly explained by order effects given a small drop in the overall standard deviation from P1 to P2.

**Table 2.**
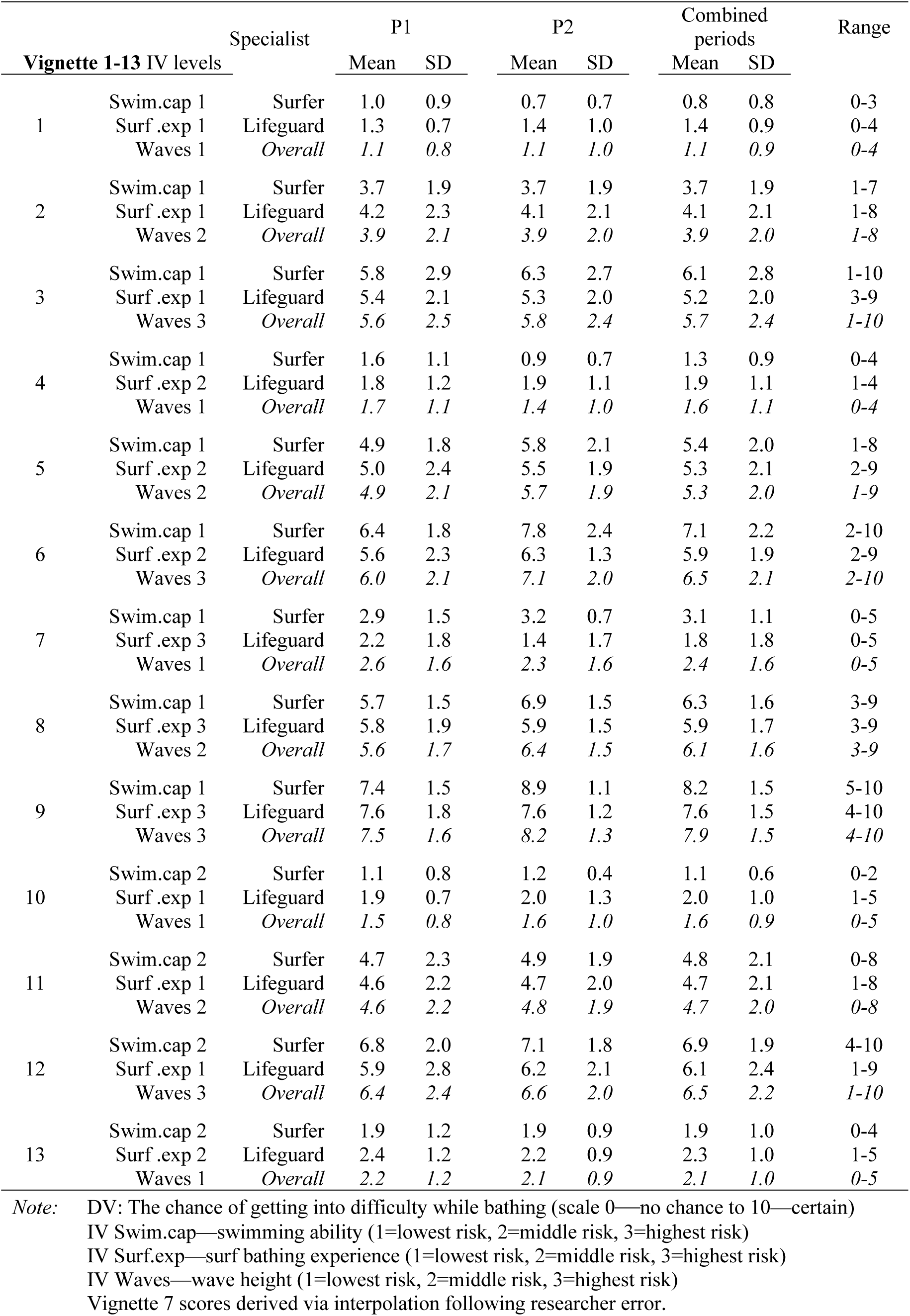
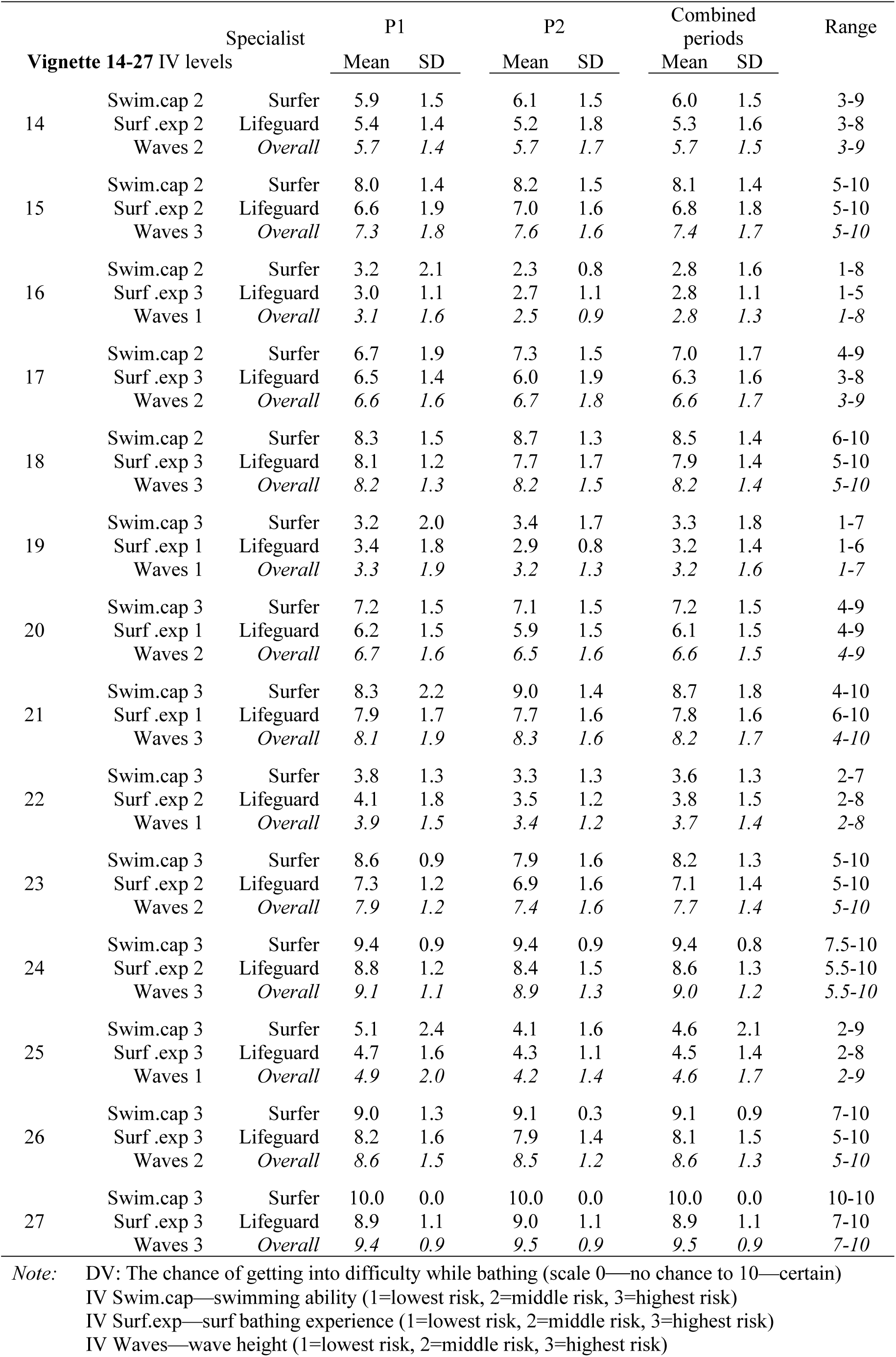
Specialist judges mean ratings of DV for vignette cells by group.

Applying the Bonferroni correction (*p*<0.002) left vignette 27 as the only statistically significant difference between specialists groups. Thus surfers’ and lifeguards’ ratings of the DV by vignette IV order levels were considered to be from the same population of specialists. Fig 2 shows the overall pattern of combined means for each of the 27 vignettes.

**Fig 2.**
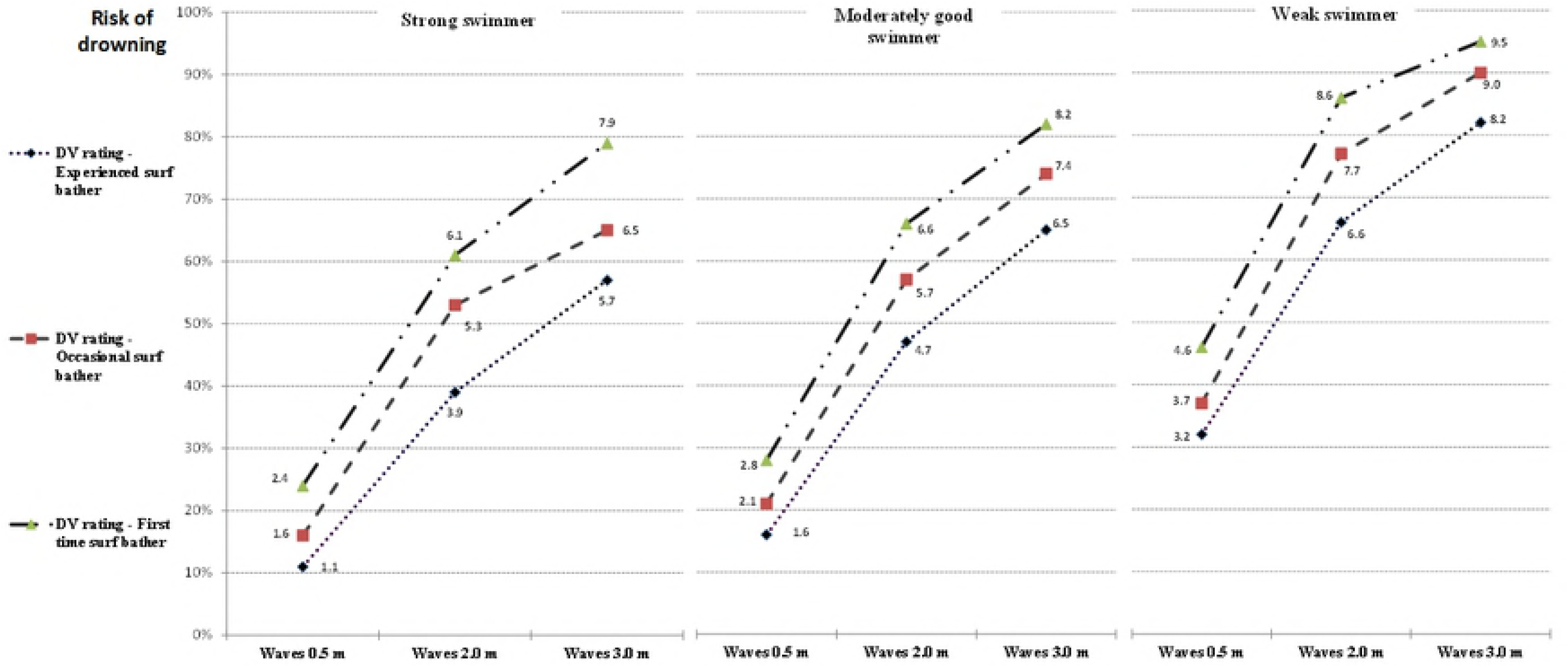
Estimated marginal mean drowning risk scores on DV for swimming ability by surf bathing experience and wave height.

The repeated-measures ANOVA procedure is robust to violations in the normality assumption of DV distributions [26]. Nevertheless, each vignette distribution for combined periods and judges was assessed for normality. Visual appearance approximated normal distributions. No distribution was significantly skewed (*p*<0.05). Four (14.8%) vignette cell distributions were significant for kurtosis (*p*<0.05) due to a high peak score (many judges chose the same rating score). Based on these results, the DV data were considered suitable for further analysis without transformation.

### Repeated-measures factorial ANOVA

A repeated-measures three-way ANOVA determined significant effects and polynomial contrasts (*p*<0.05) between the chance of getting into difficulty in the water (DV) and IVs swimming ability, surf bathing experience, and wave height, and IV interactions. Table 3 lists marginal means and standard error scores on the DV for each level of the three IVs. The overall hypothesised pattern of drowning risk posed by IV factor levels was reflected in the expert ratings.

**Table 3.**
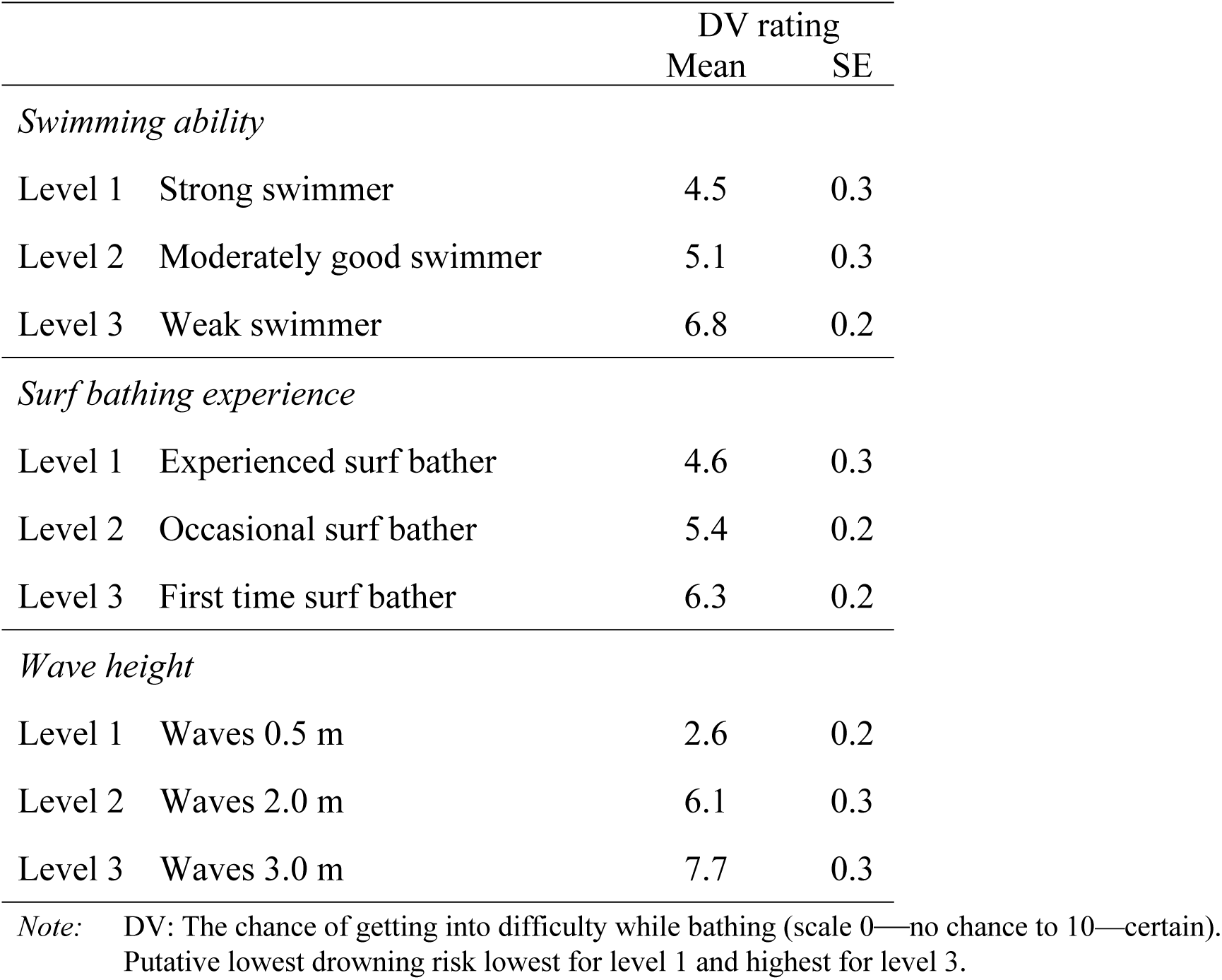
Estimated marginal means and standard errors for three drowning risk levels for IVs.

|  |  | DV rating |  |
| --- | --- | --- | --- |
|  |  | Mean | SE |
| *Swimming ability* |  |  |  |
| Level 1 | Strong swimmer | 4.5 | 0.3 |
| Level 2 | Moderately good swimmer | 5.1 | 0.3 |
| Level 3 | Weak swimmer | 6.8 | 0.2 |
| *Surf bathing experience* |  |  |  |
| Level 1 | Experienced surf bather | 4.6 | 0.3 |
| Level 2 | Occasional surf bather | 5.4 | 0.2 |
| Level 3 | First time surf bather | 6.3 | 0.2 |
| *Wave height* |  |  |  |
| Level 1 | Waves 0.5 m | 2.6 | 0.2 |
| Level 2 | Waves 2.0 m | 6.1 | 0.3 |
| Level 3 | Waves 3.0 m | 7.7 | 0.3 |

*Note:* DV: The chance of getting into difficulty while bathing (scale 0—no chance to 10—certain). Putative lowest drowning risk lowest for level 1 and highest for level 3.

### Hypotheses tests

The model resulted in simple main effects for the three IVs on the DV; swimming ability, *F*(1.21, 20.62) = 77.87, *p*<0.001, partial *η*^2^ = 0.82 (95% CI: 0.62 to 0.88), surf bathing experience, *F*(2, 34) = 99.27, *p*<0.001, partial *η*^2^ = 0.85 (95% CI: 0.74 to 0.90), and wave height, *F(*1.30, 22.05) = 227.95, *p*<0.001, partial *η*^2^ = 0.93 (95% CI: 0.85 to 0.95). All levels within each IV differed following pairwise comparisons with Bonferroni correction (*p*<.001; Table 3). Polynomial contrasts revealed a significant linear trend for the three IVs; swimming ability, *F*(1,17) = 88.51, *p*<0.001, partial *η*^2^=0.84, surf bathing experience, *F(*1,17) = 135.01, *p*<0.001, partial *η*^2^=0.89, and wave height, *F(*1,17) = 5277.04, *p*<0.001, partial *η*^2^=0.94 with significant quadratic trends for swimming ability, *F*(1,17) = 32.07, *p*<0.001, partial *η*^2^=0.65 and wave height, *F(*1,17) = 52.43, *p*<0.001, partial *η*^2^=0.76. A significant effect was found for the interaction of surf bathing experience and wave high, *F(*4, 68) = 3.03, *p*=0.02, partial *η*^2^ = 0.15 (95% CI: 0.00 to 0.27). Fig 3 shows the marginal mean scores. Polynomial contrasts found a significant linear interaction within the quadratic pattern for waves, *F(*1, 17) = 8.15, *p*=0.01, partial *η*^2^ = 0.32. A *Post-hoc* contrast comparing experienced beach swimmers to occasional beach swimmers in waves 0.5 and 2.0 m showed a significant interaction, *F*(1, 35) = 17.85, *p*<0.001, partial *η*^2^ = 0.34 (95% CI: 0.10 to 0.52).

**Fig 3.**
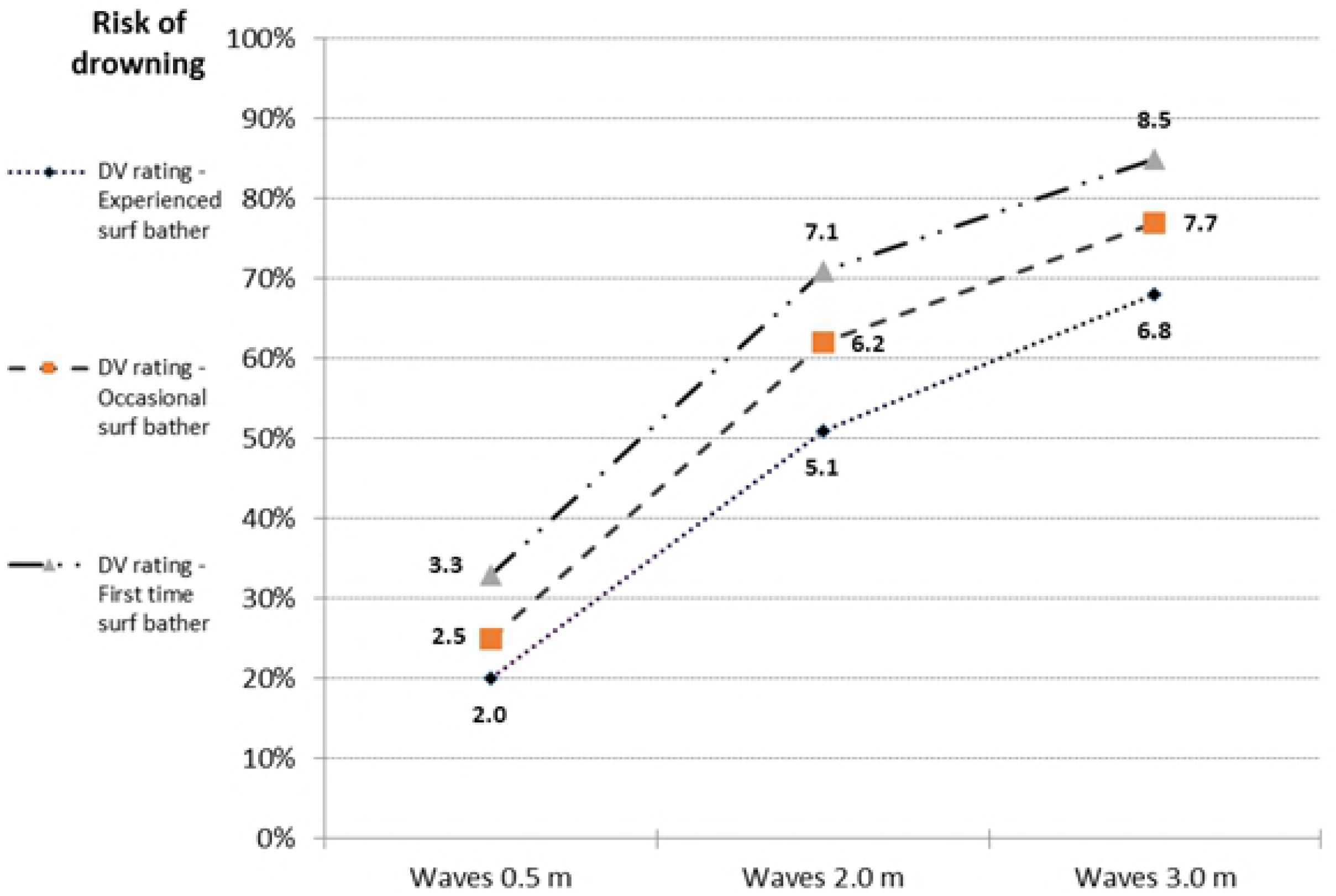
Estimated marginal mean drowning risk scores for surf bathing experience by wave height.

### Model estimation

*η*^2^ as a measure of contributed ANOVA model variance are reported in Table 4. In total, the three IVs and interactions explained 75 percent of the variability in the DV. The effects sizes are smaller and in different proportions to partial *η*^2^ due to the different base used for calculations [21].

**Table 4.**
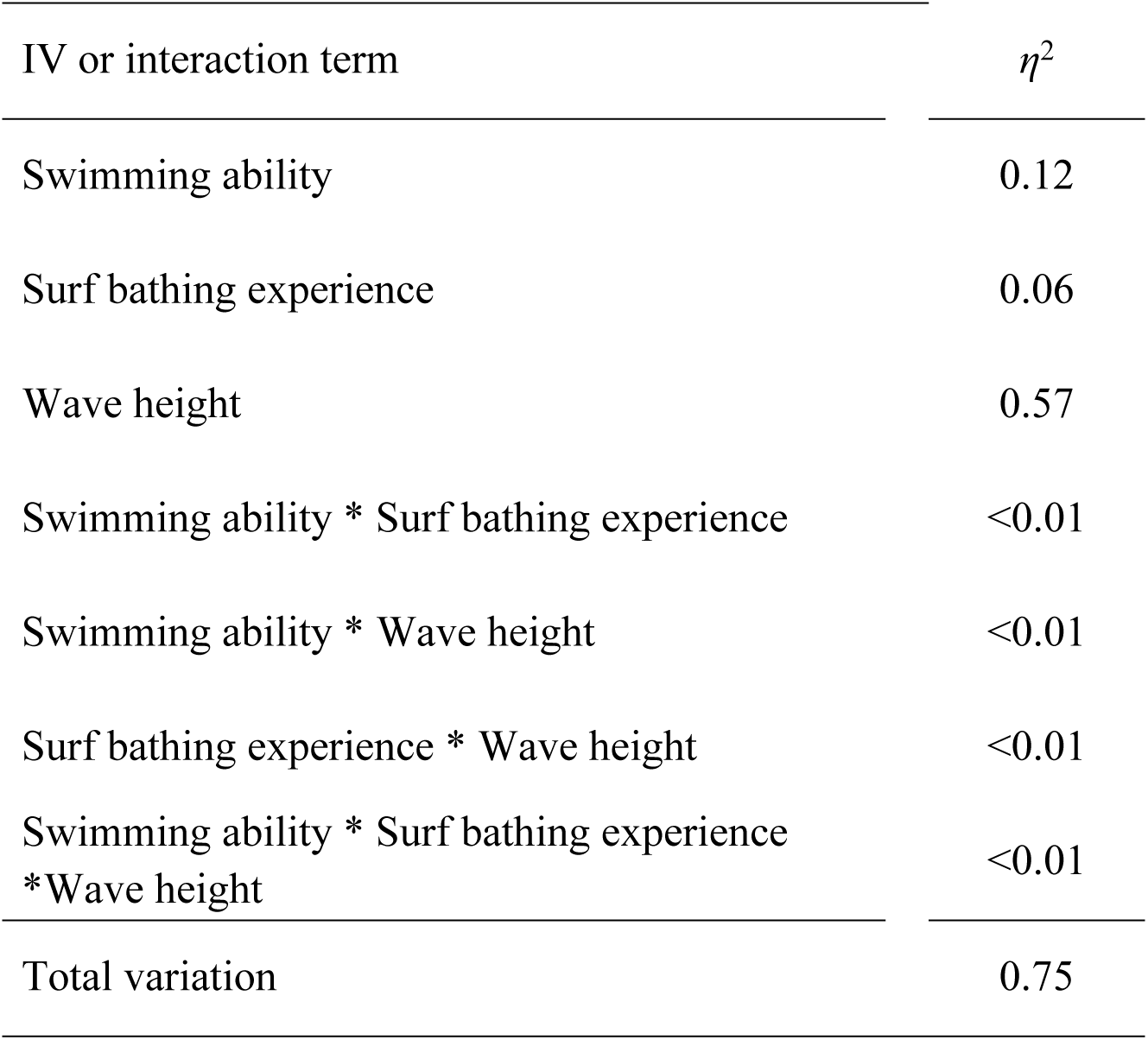
Variation in the DV explained by the IVs and interactions.

## Discussion

Eighteen lifeguards or surfers, meeting study specifications, were considered suitable judges of surf bather drowning putative risk factors based on their experience in surf activities. Each judge rated the likelihood of a person requiring rescue in 27 scenarios in a Latin square arrangement for unique combinations of three levels for swimming ability, surf bathing experience and wave height, with replication. Due to similarities of ratings, judges were considered to be from the same population and data for time-periods were pooled. This increased the study’s statistical power by reducing the proportion of error terms from presumed carryover effects.

The study found that swimming ability, surf bathing experience and wave height influenced the risk of surf bather drowning. This risk reduced when: swimming capability increased, surf bathing experience increased, or wave height decreased. The interaction between surf bathing experience and wave height suggests that drowning risk to novice surf bathers increases disproportionately at greater wave heights compared to surf bathers with more experience. No other interactions were found in the model.

The mean ratings for the 27 uniquely ordered vignettes provided a consistent pattern of rated drowning risk. This pattern *fell* as expected, imparting face validity to the study’s method and sample. DV mean cell ratings ranged from 11 percent chance of drowning for the IV combination at the lowest level of presumed risk to 95 percent chance at the highest IV risk combination. This suggests that judges considered *all* surf bathers to carry drowning risk (under scenario conditions), regardless of their skills, experience, and surf conditions.

Simple main effects of the three IVs were significant, meaning that alone each contributed variance to the DV. H1, *the three IVs will each produce an effect on the DV*, is therefore supported. Each IV accounted for a high proportion of variance within the DV, ignoring that shared with other IVs. By converting partial *η*^2^ results to percentages, these were 82, 85, and 93 percent for swimming ability surf bathing experience, and wave height respectively. Tabachnick and Fidell (p. 188) suggest that repeated-measures “produces a better guess of the effect size” compared to using a one-way ANOVA design for each factor [27]. For H2, significant linear trends for IV levels, in the expected drowning risk order, were found for each IV. The finding validated the specification of each scale as a presumed predictor of surf bather drowning risk.

Although a linear pattern of factor level distribution was strongest for each IV (based on partial *η*^2^), swimming ability and wave height also formed quadratic patterns within factors. For swimming ability, the increased drowning risk between ratings for a strong swimmer and moderately good swimmer was less pronounced relative to that between the moderately good swimmer and the weak swimmer. In contrast, the factor pattern for surf bathing experience was constant and this IV also showed less total rating variation in the DV compared to the other two IVs. Wave height showed the greatest difference in estimated marginal means from lowest to highest risk. The proportionally greater increased rating in drowning risk between 0.5 and 2 m wave height than between 2 and 3 m wave heights supports Short’s beach hazard rating system, where wave height of 0.5 m carries a beach hazard (drowning) rating of 4 (safest), 2.0 m is rated 7, and 3.0 m is rated 9 (least safe) [19].

For H3, of the three first order interactions between IVs, only that between surf bathing experience and wave height was significant, with medium effect size. Increasing wave height from 0.5 to 2.0 m presents greater proportional increase in drowning risk for occasional surf bathers compared to experienced surf bathers. The second order interaction between the three IVs was not significant. It was anticipated that drowning protection provided by strong swimming ability would increase more than proportionally at larger wave heights compared to less able swimmers. The lack of identified interactions (bar one) suggests that judges largely rated vignettes in a summative fashion based on an estimated risk contribution at the specified level for each IV.

This finding is consistent with previous studies using similar methods [14, 33]. Only one interaction being identified (assuming interactions exist) may result from judges’ failure to understand the situation correctly or the sensitivity of the research design to identify configural effects [34]. Alternatively, this result may accurately represent the judgment process, whether or not this represents the true situation [35].

Overall, this study provides *some* evidence that surf bathing specialists judge drowning risk using a configural process. Perhaps the frequent failure of previous studies to identify significant interactions between variables is explicable by the nature of the judgement task. Surf bathing experience (human-related factor) and wave height (environmental factor) are conceptually very different and dynamic variables, yet interactions would be expected. The identified interaction documents a configural judgement process used by judges to assess drowning risk from the interplay of environmental and human factors.

## Limitations

### Methodological

Although not reviewed here, statisticians debate the suitability of the long established repeated-measures factorial ANOVA where other procedures (e.g., MANOVA) may have less restrictive assumptions or may better model simple interactions [22, 36]. The repeated-measures ANOVA in this study met required assumptions and provided an appropriate test of the hypotheses. The decision to group or pool data may be challenged on strict statistical grounds. For example, a small proportion of comparisons (3.7%) identified differences in ratings between lifeguards and surfers following Bonferroni corrections. Essentially though, this limitation was counterbalanced by increased statistical power and dilution of order effects derived from combining specialist samples.

### Sample size and selection

Only 18 judges, drawn from a convenience sample, participated. Drowning risk ratings may be biased by these judges’ particular experiences and knowledge. Relevant to the study aim, the repeated-measures approach has advantage over completely randomised study designs through requiring fewer participants while having increased power and precision [26]. The consistency between drowning risk ratings from two distinct specialist groups suggests the study findings may be generalisable all surf bathing experts.

### Judges’ linear interpretation

The reality of vignette scenarios may be questioned. Judges may, for example, have perceived as contradictory or unrealistic the scenario denoting a subject as a weak swimmer with extensive surf bathing experience, a form of common method bias [37]. Such a perception could have encouraged a linear process for rating IVs and so explain the lack of interaction. At any rate, this study was limited to clearly defined circumstances. Findings restrict to the influence on drowning risk from three variables specified as fixed factors, within a general scenario specified (S2 Table), using an untested proxy measure for surf bather drowning.

### Unplanned researcher error

One IV level in a single Latin square cell was incorrect. Such data errors or losses are not unusual in studies with similar research designs where estimating scores is a satisfactory approach [22] and corrections will not have a disproportional effect on the overall results [38]. Although the method for estimating missing cell scores for individual judges was crude, results obtained by correction remained consistent with the mean DV score patterns found for each of the other 26 vignette ratings.

### Implications

Based on the study findings, surf bathers are at higher drowning risk, compared to other surf bathers, where they have inferior swimming ability, less surf bathing experience, or face larger waves. Although these findings are intuitive, in the absence of robust epidemiological data, this study provides the best available evidence supporting these conclusions. Regardless of the study limitations, comparisons between IV mean levels suggest that surf bather drowning results from a complex mix of person and situation variables and provide evidence on candidate risk factors to guide further investigation of risk and support drowning prevention strategies.

International tourists in Australia have a higher rate of surf bather drowning relative to Australian residents [6] and police reports in coronial records suggest some decedents lacked experience in surf conditions. Tourist awareness programmes on surf risk and deployment of lifeguards or surveillance drones to popular tourist areas may mitigate this risk [39-41]. Specifically, bathers with little or no surf experience should be aware that strong pool swimming ability may provide insufficient protection from drowning. Particularly for men, surf inexperience may translate to overconfidence in one’s ability to meet prevailing wave conditions [18]. Similarly, expansion of surf awareness and safety programmes (e.g. *Nippers* program for children) on surf beaches during high seasons may contribute to building surf competency and reducing over-confidence [42-43].

Additionally, meteorological reports of surf conditions could incorporate indications of risk level (e.g., not suitable for inexperienced surf swimmers), somewhat similar to current ultraviolet radiation warnings [44]. In Australia, drowning risk indicators may be integrated within the Beachsafe website that provides detailed bather-related information for surf beaches [45]. Technical advances in inflatable lifejackets, originally designed for big wave surfers, may also offer drowning protection, especially for weak swimmers or inexperienced surf bathers [46]. Waterproof GPS tracking devices may also potentially aid in bather surveillance and timely rescue [47]. These and other possible countermeasures require careful evaluation before their efficacy in drowning prevention is assumed.

With regard to future epidemiological observations studies, analysis of expert judgments provides a step towards distinguishing the roles of key variables within this complexity. The extent to which the findings represent the true surf drowning situation requires comparison to a gold standard gained only through rigorous epidemiological designs. The availability of such evidence would be ideal, but meanwhile, the method reported here provides a useful alternative or preliminary investigation of injury risk factors to reduce drowning.

## Conclusion

The proof of method reported here offers an important avenue for investigating significant health problems, including unintentional injury, by providing a *window* into the true situation. Expert judgement carefully collected and analysed, can be used to document and assess the roles of risk contributions from putative causal factors that determine health outcomes. This method provides injury researchers with a rapid low cost tool for data collection comparable to that obtained through resource intensive epidemiological designs. A method based on expert judgement of course cannot replace these more robust designs, but may prove a useful substitute or preliminary method for generating new knowledge to address health problems and improve outcomes.

## Acknowledgments

The authors express their thanks to the surf specialists who provided their time for participation in the research study.

## Supporting information

**S1 Table Scale Specification**

**S2 Table Questionnaire instructions**

